# Sexually dimorphic effects of prenatal alcohol exposure on the murine skeleton

**DOI:** 10.1101/2024.03.27.586945

**Authors:** Lucie E Bourne, Soher N Jayash, Lysanne V Michels, Mark Hopkinson, Fergus M Guppy, Claire E Clarkin, Paul Gard, Nigel Brissett, Katherine A Staines

**Author notes:** Email addresses: LEB; SNJ; LVM; MH; FMG; CEC; PG; NB; KAS. **Corresponding author** Katherine Staines School of Applied Sciences, University of Brighton, Lewes Road, Brighton BN2 4GJ.

## Abstract

**Background:** Prenatal alcohol exposure (PAE) can result in lifelong disabilities known as foetal alcohol spectrum disorder (FASD) and is associated with childhood growth deficiencies and increased bone fracture risk. However, the effects of PAE on the adult skeleton remain unclear and any potential sexual dimorphism is undetermined. Therefore, we utilised a murine model to examine sex differences with PAE on *in vitro* bone formation, and in the juvenile and adult skeleton.

**Methods:** Pregnant C57BL/6J female mice received 5% ethanol in their drinking water during gestation. Primary calvarial osteoblasts were isolated from neonatal offspring and mineralised bone nodule formation and gene expression assessed. Skeletal phenotyping of 4- and 12-week-old male and female offspring was conducted by micro-computed tomography (µCT), 3-point bending, growth plate analyses, and histology.

**Results:** Osteoblasts from male and female PAE mice displayed reduced bone formation, compared to control (≤30%). *Bglap* and *Ahsg* were upregulated with PAE in both sexes compared to control, whereas *Vegfa*, *Bmp6*, *Tgfbr1* and *Flt1* were downregulated in PAE male osteoblasts only. In 12-week-old mice, µCT analysis revealed a sex and exposure interaction across several trabecular bone parameters. PAE was detrimental to the trabecular compartment in male mice compared to control, yet PAE females were unaffected. Both male and female mice had significant reductions in cortical parameters with PAE. Whilst male mice were negatively affected along the tibia length, females were only distally affected. Posterior cortical porosity was increased in PAE females only. Mechanical testing revealed PAE males had significantly reduced bone stiffness compared to controls; maximum load and yield was reduced in both sexes. PAE had no effect on total body weight or tibial bone length in either sex. However, total growth plate width in male PAE mice compared to control was reduced, whilst female PAE mice were unaffected. 4-week-old mice did not display the altered skeletal phenotype with PAE observed in 12-week-old animals.

**Conclusions:** Evidence herein suggests for the first time that PAE exerts divergent sex effects on the skeleton, possibly influenced by underlying sex specific transcriptional mechanisms of osteoblasts. Establishing these sex differences will support future policies and clinical management of FASD.

**Plain English summary:** Prenatal alcohol exposure (PAE) can lead to a set of lifelong cognitive, behavioural, and physical disabilities known as foetal alcohol spectrum disorder (FASD). FASD is a significant burden on healthcare, justice and education systems, which is set to worsen with rising alcohol consumption rates. FASD children have an increased risk of long bone fracture and adolescents are smaller in stature. However, sex differences and the long-term effects of PAE on the skeleton have not been investigated and was the aim of this study. Using a mouse model of PAE, we examined the function and gene expression of bone-forming cells (osteoblasts). We then analysed the skeletons of male and female mice at 12-weeks-old (adult) and 4-weeks-old (juvenile). PAE reduced osteoblast bone formation in both sexes, compared to control. Differential gene expression was predominantly observed in PAE males and largely involved genes related to blood vessel formation. High resolution x-ray imaging (micro-CT) revealed PAE had a detrimental effect on the inner trabecular bone component in 12-week-old male mice only. Analysis of the outer cortical bone revealed that whilst both male and female PAE mice were negatively affected, anatomical variations were observed. Mechanical testing also revealed differences in bone strength in PAE mice, compared to control. Interestingly, 4-week-old mice did not possess these sex differences observed in our PAE model at 12 weeks of age. Our data suggest PAE has detrimental and yet sex-dependent effects on the skeleton. Establishing these sex differences will support future policies and clinical management of FASD.

**Highlights:**

- Primary calvarial osteoblasts isolated from male and female PAE mice displayed reduced mineralised bone nodule formation and differential gene expression compared to control.
- PAE had a detrimental effect on trabecular bone parameters in 12-week-old male mice only.
- PAE leads to spatial variation in cortical bone parameters and geometry, with male mice negatively affected along the tibia length and female mice only affected at the distal end.
- Mechanical testing revealed PAE male mice had significantly reduced bone stiffness compared to controls; PAE in both sexes reduced maximum load and yield.
- 4-week-old mice did not display the altered skeletal phenotype with PAE observed in 12-week-old animals.

## Background

There is increasing evidence to suggest that the intrauterine environment influences lifelong skeletal health [1–3]. Prenatal alcohol exposure (PAE) can lead to lifelong disabilities known as Foetal Alcohol Spectrum Disorder (FASD). FASD is associated with over 400 comorbidities and adverse outcomes in later life [4, 5]. It is characterised by a variety of cognitive and physical impairments, notably its effects on brain development, facial dysmorphology, central nervous system (CNS) malformations, and has since been associated with learning difficulties [6]. It is a leading cause of developmental disability, affecting approximately 1-10% of children in the USA and Europe, and resulting in a significant burden on healthcare, social, justice and education systems [7–9]. Recent analysis of the Avon Longitudinal Study of Parents and Children estimated that the screen prevalence in this UK population-based cohort was 17% [10], whilst global prevalence of children born with foetal alcohol syndrome (FAS), the most severe form of FASD, is estimated to be >100,000 per year [11].

Whilst the effects of PAE on craniofacial differences are well documented [12–15], studies investigating the clinical effects on the rest of the skeleton are limited. Early small-scale studies and case reports highlighted skeletal malformations including digital abnormalities (e.g. clinodactyly), carpal fusion, radioulnar synostosis, cervical spine abnormalities, and skeletal dysplasia [16–18]. Since then, growth deficiencies have become a hallmark of FASD and are observed in children exposed to alcohol *in utero*, at any point during pregnancy [13, 19, 20]. Additionally, there is increasing evidence that adolescents with FASD are shorter, have a reduced head circumference and a delayed mean bone age [19, 21–23]. Similarly, PAE is associated with an increased risk of a long bone fracture in childhood [24], and with reduced bone mineral density (BMD) and lean tissue mass in adolescents [23].

In animal models, there is further evidence that PAE results in skeletal irregularities. Skeletal malformations such as fused or abnormal ribs, delayed hindlimb ossification, forelimb anomalies and digit defects have been observed in the offspring of mice given ethanol via intraperitoneal injection or as part of their diet [25, 26]. Foetal growth, bone development and percent ossification are also disrupted in rat foetuses during high and moderate ethanol exposure [27–29], alongside disturbances to calcium homeostasis [27, 30]. Similarly, studies in sheep show high dose alcohol exposure during pregnancy reduces foetal bone length and strength [31–33]. Further, alongside its effects on bone, PAE also alters cartilage development as it disrupts the organisational zones of the growth plate, decreasing the length of the resting zone and increasing the length of the hypertrophic zones of chondrocytes [34].

Whilst these data suggest a detrimental effect of PAE on bone development, the long-term effects on the adult skeleton have not been fully investigated. Similarly, whilst sex differences in human FASD populations have been observed across various systems and clinical outcomes, including neurodevelopment, cognitive performance, endocrine disorders, facial dysmorphology, pre- and post-partum mortality, body composition and metabolic abnormalities [35–37], any potential sexual dimorphism associated with bone architecture and composition is unclear [23]. As evidence suggests that diagnosis and disease presentation of these patients can alter between males and females, establishing these sex differences will support future policies and clinical management [35]. In addition, addressing sex bias in skeletal research, particularly in preclinical models, is necessary for effective translation into a clinical setting [38]. Therefore, in this study we utilised a murine model to examine potential sex differences with PAE on *in vitro* bone formation and in the juvenile and skeletally mature skeleton.

## Methods

### Animals

Pregnant C57Bl/6J females received 5% ethanol in their drinking water during gestation. Fluid intake between control (water-only) and alcohol-exposed females was unchanged. All mice utilised in these studies were kept in controlled conditions at the University of Brighton. All tissue isolation and experimental procedures were performed in accordance with the UK Animals (Scientific Procedures) Act of 1986 and regulations set by the UK Home Office and local institutional guidelines. Tissues were collected from male and female mice at 4- and 12-weeks of age, following euthanasia by cervical dislocation. For 12-week animals: n=8 control males, n=11 PAE males, n=9 control females, n=8 PAE females. For 4-week animals: n=5 control males, n=9 PAE males, n=10 control females, n=7 PAE females. Analyses were conducted blindly to minimise the effects of subjective bias. Animal studies were conducted and reported in line with the ARRIVE guidelines.

### Primary osteoblast cultures

Osteoblasts were isolated from the calvariae of male and female 3–5-day old mice from control and PAE dams by trypsin/collagenase digestion, as previously described [39, 40]. Timed matings were performed to ensure, where possible, cells were isolated from control and PAE offspring at the same time. Cells were expanded in Alpha Minimum Essential Medium supplemented with 10% foetal bovine serum (FBS), 100 U/ml penicillin, and 100μg/ml streptomycin (mixture abbreviated to αMEM). Following plating in 6-well plates at 10^5^ cells/well and 12-well plates at 5×10^4^ cells/well, osteoblasts were cultured for up to 21 days in αMEM supplemented with 2mM β-glycerophosphate and 50μg/ml ascorbic acid, with half medium changes every 3 days. Time points in osteoblast cultures were: differentiating (Day 7), mature (Day 14), and mature, bone forming (Day 21). All experiments were performed on cells that were isolated, expanded and plated; the cells were not passaged at any stage.

### Assessment of mineralisation

Experiments were terminated after 21 days by fixing the cells in 2.5% glutaraldehyde for 5 min. Plates were imaged at 800dpi using a flat-bed scanner and the total area of bone nodules was quantified by image analysis, as described previously [40]. Alizarin red (1% *w/v*) stained images were obtained, as previously described [41].

### Quantitative polymerase chain reaction (qPCR) and PCR array

RNA was isolated and purified from primary osteoblast cultures at day 7 and 14 of culture using the Qiagen RNeasy mini kit, according to the manufacturer’s instructions. For the PCR array, 2µg of RNA from day 7 cultures was reverse transcribed using a Qiagen RT^2^ First Strand Kit, according to manufacturer’s instructions. Gene expression changes associated with osteogenesis were assessed using an RT^2^ Profiler™ PCR Array (PAMM-026Z, Qiagen). A fold regulation threshold of 2 was used to determine up- or down-regulation of each gene in male and female PAE mice compared to control and data is presented as log_2_ fold change, relative to the control. Subsequently, qPCR analysis of key genes identified in the PCR array was conducted at days 7 and 14 of culture. RNA (2µg) was reverse transcribed using a Qiagen QuantiTect RT kit, according to the manufacturer’s instructions. The PCR was performed using cDNA (50ng) and a Brilliant II sybr green mastermix (Agilent). Both the PCR array and qPCR were run on an Agilent AriaMx PCR machine. Gene expression data for the qPCR were normalized to β-actin and analysed using the ΔΔCt method [42]. Primer sequences can be found in Suppl. Table 1.

### MicroCT (µCT) analysis

Scans of the left tibia were performed with an 1172 X-Ray microtomograph (Skyscan). High-resolution scans with voxel size of 5µm were acquired (50kV, 200µA, 0.5mm aluminium filter, 0.6° rotation angle). The projection images were reconstructed using NRecon software (Skyscan). Each dataset was rotated in DataViewer (Skyskan) to ensure similar orientation and alignment for analysis. Segmentation of the datasets was performed in CTAn software (Skyscan) to exclude the fibula. 3D analysis of the trabecular bone was performed using a volume of interest of 5% of the total bone length. This region extended distally from the bottom of the growth plate where the primary spongiosa is no longer visible towards the diaphysis. A binarised image with a minimum threshold set to 85 was used across all datasets and 3D measurements taken in CTAn. BMD measurements were calibrated using the same measurements with appropriate phantoms. Cortical bone was analysed using a 2D whole-bone approach to visualise changes across the entire tibia [43]. To ensure trabecular bone was not included in the analysis, the proximal and distal 10% of the tibial length were excluded. Slice by slice measurements were created in CTAn using binarised images with a minimum threshold set to 80. 3D images for cortical and trabecular regions were created using Avizo software (ThermoFisher Scientific).

### Cortical Porosity

Using the tibial µCT scans, 200 slices starting at the tibiofibular junction were imported into FIJI ImageJ. First, thresholds were determined to create binary images, followed by ‘Keep Largest Region’ to isolate the tibia. Using ‘Moment of Inertia’, unaligned scans were aligned so each region would fall in a quadrant. Anterior, medial, lateral, and posterior regions were expected to show spatial differences [44] and therefore extracted by halving the canvas size. A filled mask was created and the marrow cavity subtracted, resulting in a cortical tissue mask. Intracortical canals were extracted by subtracting the binary tibial image from this mask, with no extraction of osteocyte lacunae due to low resolution. With BoneJ’s ‘Particle Analyser’, cortical porosity in % (canal volume normalised for cortical tissue volume) was assessed [45]. 3D reconstructions were created in ORS Dragonfly using ‘Contour Mesh’ for visualisation.

### Three-point bending

Left femurs were cleaned of soft tissue and frozen in distilled water at −20°C. Three-point bending was carried out on thawed bones using a Lloyd LRX5 materials testing machine (Lloyd Instruments) fitted with a 100N load cell. The span was fixed at 10mm and bones loaded with a crosshead speed of 1mm/min until failure, as previously described [46]. Load extension curves were generated to identify maximum load. Stiffness and yield were calculated from the slope of the linear region of the load extension curve using polynomial fit.

### Histological analysis

Right tibiae were fixed for 24 hours in 4% paraformaldehyde before being decalcified in 10% ethylenediaminetetraacetic acid (EDTA) for 21 days at 4°C, wax-embedded and 6μm coronal sections cut. Slides were stained with H&E using standard procedures. For TRAP staining 70mg napthol AS-TR phosphate (Sigma) was dissolved in 250µl N-N dimethyl formamide (Sigma) and added to 50ml 0.2M sodium acetate buffer pH 5.2. 115mg sodium tartrate dihydrate (Sigma) and 70mg fast red salt TR (Sigma) was dissolved into this solution and slides were incubated at 37°C for 2 hours. Sections were counterstained in Meyer’s haematoxylin (Sigma), washed in distilled water and mounted in aqueous mounting medium (Vector Labs) [47].

### Growth plate zone and bridge analysis

The width of the growth plate proliferating and hypertrophic zones, as well as the total growth plate width, were measured at 10 different points along the length of the growth plate based on established cell morphology. Measurements were conducted in 6µm coronal sections from the middle region of the tibia in similar location of four individual animals per experimental group, using a light microscope and ImageJ software. Growth plate bridge analysis was conducted using a 3D µCT quantification method as previously described [48]. Briefly, µCT scans of the tibiae were segmented using a region-growing algorithm within the Avizo software. The central points of all bony bridges were identified and projected on the tibial joint surface to allow spatial distribution.

### Statistical analysis

Analyses were performed using GraphPad Prism software (version 10.1.2). For *in vitro* analyses, results are presented as bar graphs with points from individual experiments (n=4 biological replicates); containing 3-6 technical replicates. For *ex vivo* skeletal phenotyping, data are presented as bar graphs with points for individual animals. Normal distribution of data was assessed using the Shapiro-Wilk normality test. For comparison between control and PAE datasets in both males and females, a two-way ANOVA followed by Tukey’s post hoc analysis was used to test for significance. qPCR data was analysed using an unpaired, two-tailed t test. Results are presented as the mean ± standard error of the mean (SEM).

For the cortical bone analysis, R software (version 4.2.2) was used to generate line graphs and perform statistical analyses. Normal distribution of data was assessed using the Shapiro-Wilk normality test. For comparison between control and PAE datasets in both males and females, a two-way ANOVA followed by Tukey’s post hoc analysis was used to test for significance. Lines represent mean, with shading representing ± SEM. Colour heatmaps display the statistical significance at matched locations along the graph showing the PAE effect (in males and females) and the sex effect (in controls and PAE). Red = *p*<0.001, green = *p*<0.01, yellow = *p*<0.05, blue = *p*>0.05 (not significant).

## Results

Utilising a murine model of PAE, the aim of this study was to examine the role PAE has on *in vitro* bone formation and skeletally phenotype mice at 4- and 12-weeks of age. Herein, osteoblasts derived from control and PAE neonates of both sexes were used to investigate the effects of PAE on bone cell function. Tibias and femurs from both male and female mice were used to investigate growth plate, trabecular and cortical bone architecture, alongside mechanical testing. Figure 1 shows a schematic of the study protocol.

**Figure 1.**
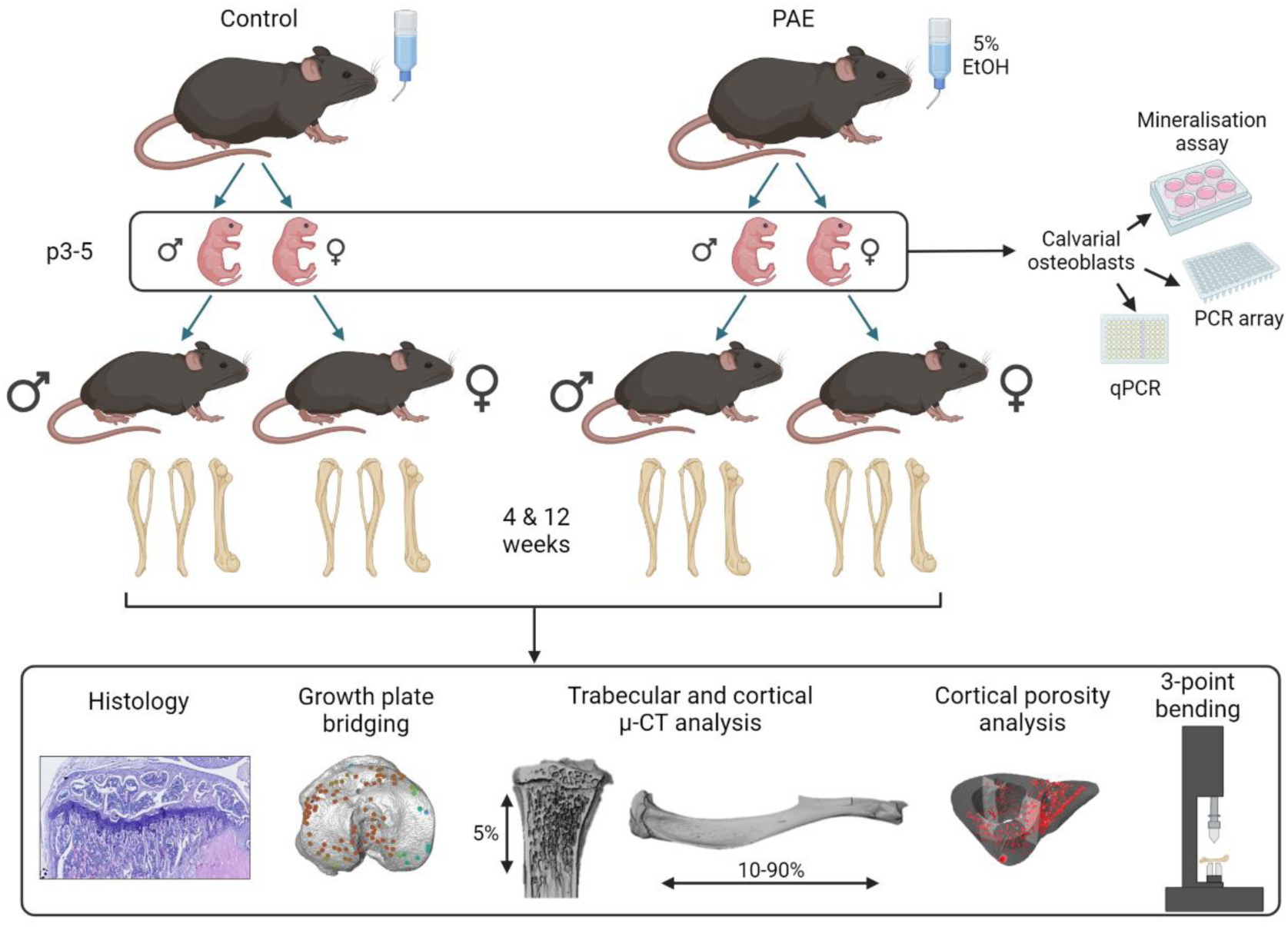
Schematic of experimental design. Pregnant dams were exposed to 5% ethanol in their drinking water (prenatal alcohol exposure (PAE)) or normal drinking water. Offspring were sacrificed at postnatal (p) days 3-5 for primary osteoblast cultures, or 4- and 12-weeks of age for skeletal phenotyping. Created with Biorender.com.

### PAE reduces in vitro bone formation in males and females but results in sex-dependent differences in osteoblast gene expression

To understand whether PAE alters osteoblast differentiation and function, primary osteoblasts derived from calvarial neonates were used as an effective model of *in vitro* bone formation [39]. Mineralised nodule formation in cells derived from both PAE males and females was reduced by ≤30%, compared to control cells (*p*<0.001; Fig. 2A & B). An osteogenesis PCR array performed at day 7 of culture, during the differentiation stage of osteoblasts, highlighted several genes that were differentially expressed between male and female osteoblasts. *Bglap* (Osteocalcin), *Csf2* and *Gli1* were upregulated in control females compared to male, whilst *Bmp6*, *Col141a* and *Flt1* were downregulated. Compared to PAE males, female PAE osteoblasts only showed an increase in *Ahsg* (Fetuin A). PAE also differentially affected osteoblast gene expression compared to control within each sex. Upregulation of genes for *Bglap* and *Ahsg* were observed in both PAE males and females compared to control, whilst *Vegfa* was downregulated in PAE from both sexes. *Bmp6*, *Tgfbr1* and *Flt1* (VEGFR) were downregulated in PAE male mice only compared to control (Table 1).

**Figure 2.**
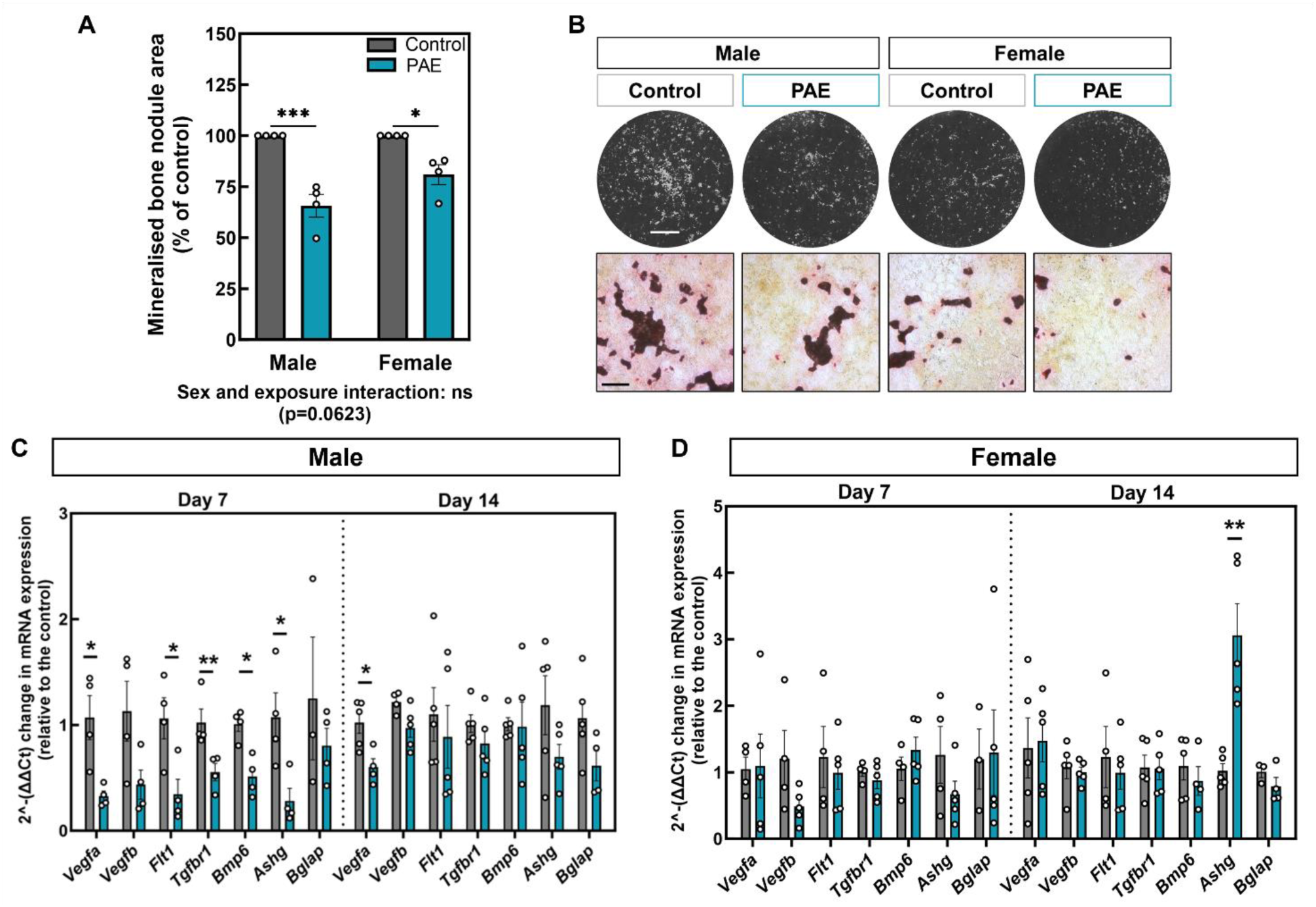
PAE reduces in vitro bone formation and results in sex-dependent differences in osteoblast gene expression. (**A**) Mineralised bone nodule area of primary calvarial osteoblasts from PAE male and female mice as a percentage of the control cultures. (**B**) Representative whole-well scans and light microscopy images of mineralised nodules. (**C**) RT-qPCR of key genes identified by RT^2^-profiler analysis (Vegfa, Vegfb, Flt1, Tgfbr1, Bmp6, Ashg, and Bglap) on days 7 and 14 of culture from osteoblasts derived from male control and PAE, and (**D**) female control and PAE mice (data normalised to β-actin). Data are presented as mean ± SEM with points showing individual experiments. *= p<0.05, **= p<0.01, ***= p<0.001. Scale bars: whole well = 0.5cm, microscopy = 200μm.

**Table 1.** Osteogenesis PCR array of control and PAE primary osteoblasts from male and female mice. Data is presented as log_2_ fold change, relative to males or relative to the control.

| Log<sub>2</sub> fold regulation |  |  |  |  |  |  |  |
| --- | --- | --- | --- | --- | --- | --- | --- |
| Gene | Control<br>Female vs<br>Male | Gene | PAE<br>Female vs<br>Male | Gene | Male<br>PAE vs<br>Control | Gene | Female<br>PAE vs<br>Control |
| *Bglap* | 1.22 | *Ahsg* | 2.08 | *Ahsg* | 2.32 | *Ahsg* | 3.78 |
| *Csf2* | 2.72 |  |  | *Bglap* | 1.94 | *Bglap* | 1.19 |
| *Gli1* | 1.02 |  |  | *Csf2* | 1.62 | *Col14a1* | 1.34 |
| *Bmp6* | -1.03 |  |  | *Mmp10* | 1.5 | *Comp* | 1.52 |
| *Col4a1* | -1.51 |  |  | *Mmp8* | 1.49 | *Mmp10* | 1.16 |
| *Flt1* | -1.36 |  |  | *Bmp6* | -1.29 | *Col2a1* | -1.25 |
|  |  |  |  | *Col4a1* | -1.12 | *Ihh* | -1.47 |
|  |  |  |  | *Flt1* | -2.64 | *Vegfa* | -1.84 |
|  |  |  |  | *Tgfbr1* | -1.03 |  |  |
|  |  |  |  | *Vegfa* | -1.84 |  |  |
|  |  |  |  | *Vegfb* | -1.12 |  |  |

To confirm the results seen in the array and assess how gene expression changes during maturation, qPCR was performed on osteoblasts derived from PAE and control neonates at both day 7 (differentiating) and day 14 (mature, mineral-forming) of culture. *Vegfa*, *Flt1*, *Bmp6* and *Tgfbr1* were all significantly downregulated in PAE males at day 7, compared to control (*p*<0.05; Fig. 2C), with a trend towards downregulation of *Vegfb* that did not reach significance (Fig. 2C). By day 14, only *Vegfa* remained downregulated in PAE males compared to control (*p*<0.05; Fig. 2C). In females, no differences were observed between control and PAE at day 7 and only an increase in *Ashg* in PAE cells was observed at day 14 (*p*<0.01; Fig. 2D).

### PAE has a detrimental effect on trabecular bone parameters in male mice, whilst female mice are protected

We next sought to examine whether the alterations in bone formation and osteoblast commitment in cells from neonatal PAE mice had lasting effects on the adult skeleton. In 12-week-old mice, µCT analysis revealed a sex and exposure interaction across a number of trabecular parameters in the tibia (Fig. 3A-I). Males and females were significantly different from each other in both control and PAE animals (*p*<0.05). However, in male mice compared to control, PAE was detrimental to the trabecular compartment. Specifically, PAE significantly reduced many of these parameters including tissue volume (*p*<0.0001; Fig. 3A), bone volume (*p*<0.0001; Fig. 3B), bone volume/tissue volume (*p*<0.05; Fig. 3C), bone surface (*p*<0.0001; Fig. 3D) and intersection surface (*p*<0.0001; Fig. 3E). The structure and architecture of the trabeculae was also altered in PAE male mice (Fig. 3F-I). Trabecular number (*p*=0.051; Fig. 3F), fractal dimension (*p*<0.05; Fig. 3G) and connectivity density (*p*<0.001; Fig. 3H) were decreased, whilst trabecular pattern factor (*p*<0.05; Fig. 3I) was increased, indicating a more disconnected and less complex trabecular compartment. However, female mice were unaffected by PAE, with no significant differences observed in any of the trabecular bone parameters examined (Fig. 3A-I). Representative 3D-reconstructed images of the trabecular compartment highlight these differences between males and females (Fig. 3J). Additional parameters and parameters that were unaffected by PAE in either sex, such as trabecular thickness, trabecular separation and bone mineral density are shown in Suppl. Fig 1.

**Figure 3.**
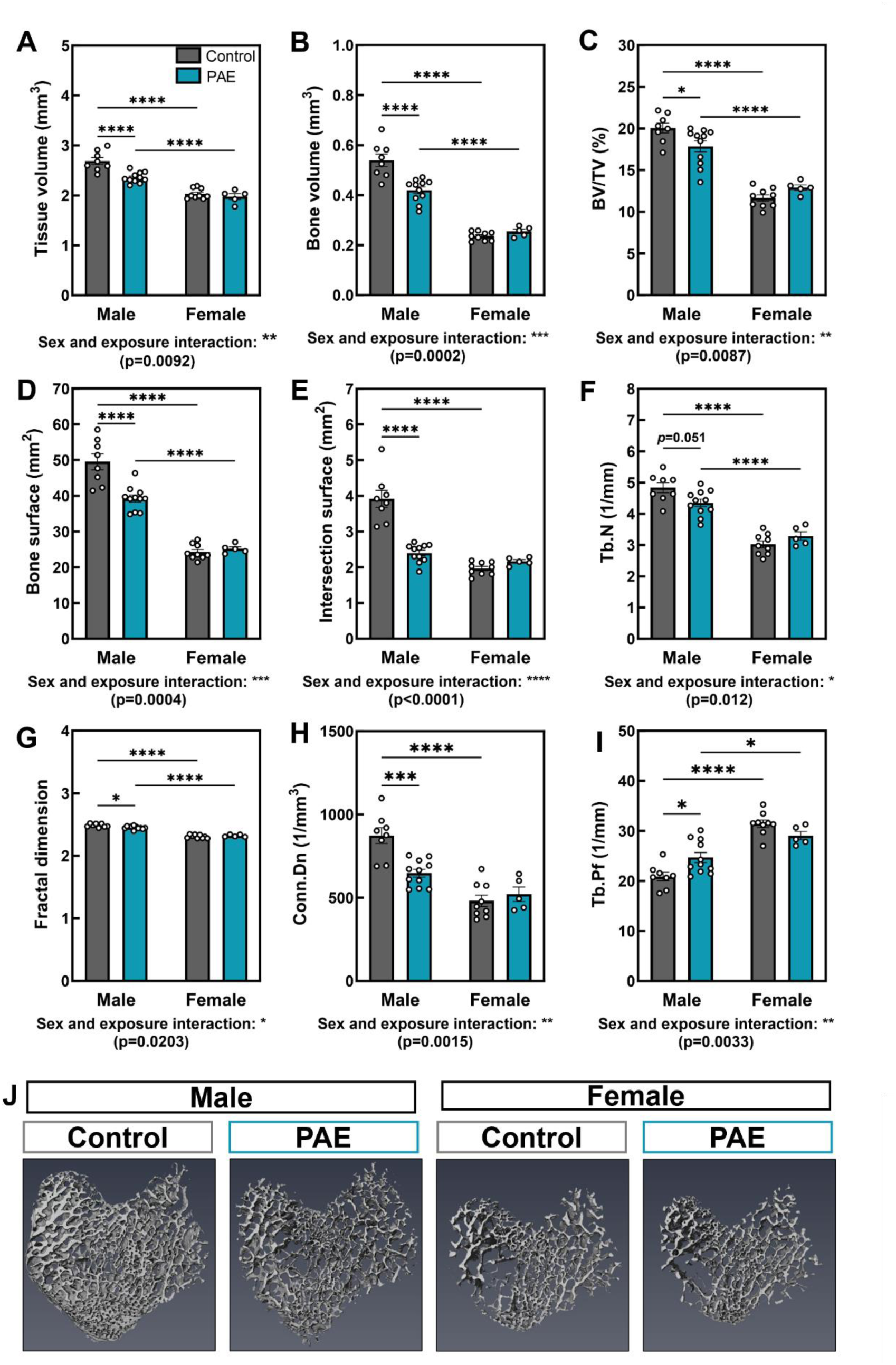
PAE has a detrimental effect on the trabecular microarchitecture in male mice. µCT analysis of (**A**) tissue volume, (**B**) bone volume, (**C**) bone volume to tissue volume ratio (BV/TV), (**D**) bone surface, (**E**) intersection surface, (**F**) trabecular number (Tb.N), (**G**) fractal dimension, (**H**) connectivity density (Conn.Dn) and (**I**) trabecular pattern factor (Tb.Pf) in male and female PAE and control mice. (**J**) Representative 3D-reconstructed images of the trabecular region in control and PAE mice. Data are presented as mean ± SEM with points showing individual animals. *= p<0.05, ***= p<0.001, ****= p<0.0001.

### PAE leads to spatial variation in cortical bone parameters, geometry and biomechanical properties in both males and females

To assess the effect of PAE along the full length of the tibia, 2D whole bone analysis was performed (Fig. 4). In both control and PAE mice, females showed a reduction in each parameter compared to males along almost the entirety of the bone (*p*<0.05; Fig 4A-F). However, in both sexes compared to control, PAE negatively affected cortical parameters including tissue area (*p*<0.05; Fig. 4A), tissue perimeter (*p*<0.05; Fig. 4B), and cortical bone area (*p*<0.05; Fig. 4C). Whilst PAE males compared to control males were affected along the entire length of the tibia, female PAE mice only showed differences to control at the distal end (Fig. 4A–C). In addition, PAE also similarly altered parameters associated with cortical bone shape in these mice, including predicted resistance to torsion *(J; p*<0.05; Fig. 4D), maximum (*I_max_*; *p*<0.05; Fig. 4E) and minimum *(I_min_*; *p*<0.05; Fig. 4F) moments of inertia. Again, these alterations were found along the length of the tibia in PAE males relative to control males, whereas PAE females were only affected more distally (Fig. 4D-F). Interestingly, across several parameters, a specific area around the tibiofibular junction (60%) was differentially affected between males and females, whereby PAE females showed reductions in these parameters compared to controls (*p*<0.01), whilst PAE males were unaffected.

**Figure 4.**
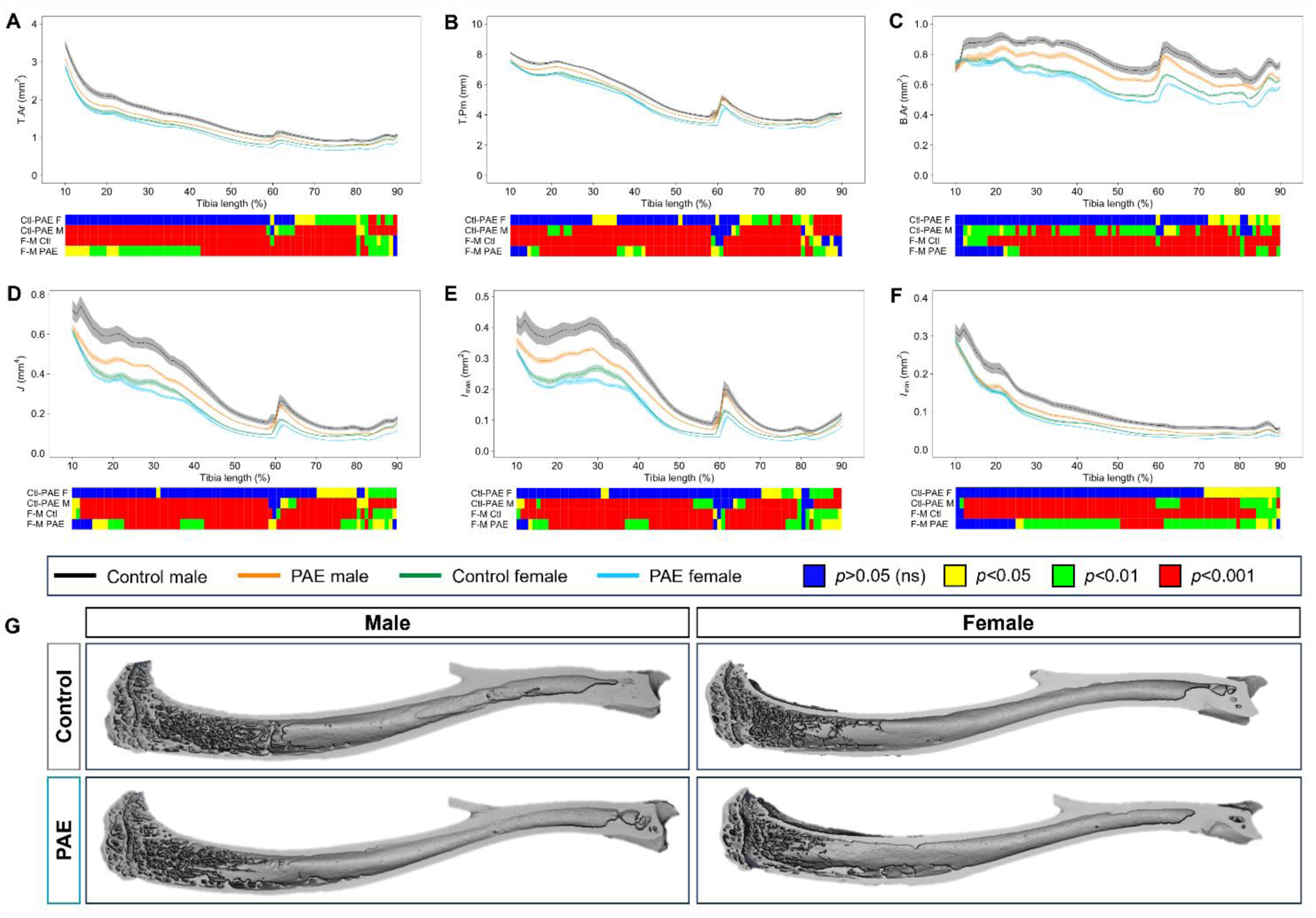
PAE leads to spatial variation in changes in tibial cortical parameters and geometry. Measurement and statistical analysis heat map for (**A**) tissue area (T. Ar), (**B**) tissue perimeter (T.Pm), (**C**) Bone area (B.Ar), (**D**) resistance to torsion (J), (**E**) maximum second moments of inertia (I_max_) and (**F**) minimum second moments of inertia (I_min_) in male and female PAE and control mice. Line graphs represent mean ±SEM for male control (black), male PAE (orange), female control (green) and female PAE (blue) mice. Graphical heat map summarises statistical differences at specific matched locations along the tibial length (10% to 90%), for the PAE effect in females (Ctl-PAE F) and males (Ctl-PAE M), and the sex effect in control (F-M Ctl) and PAE (F-M PAE) groups. Red = p<0.001, green = p<0.01, yellow = p<0.05, blue = p>0.05 (not significant).

We next sought to examine whether the cortical changes observed were associated with any changes in cortical porosity, using the tibiofibular junction as an anatomical landmark (Fig. 5A). Whilst there were no differences in total cross-sectional cortical porosity between control and PAE mice in both sexes (Fig. 5B), regional analysis revealed changes in the posterior region of the tibia in female PAE mice only. In both sexes, the posterior region contained greater porosity compared to the anterior, lateral and medial regions (*p*<0.05; Fig. 5C). However, PAE females had greater porosity in this region compared to control females (*p*<0.001) and to PAE males (*p*<0.01). PAE males were unaffected compared to control and no sex differences were observed in control animals (Fig. 5C).

**Figure 5.**
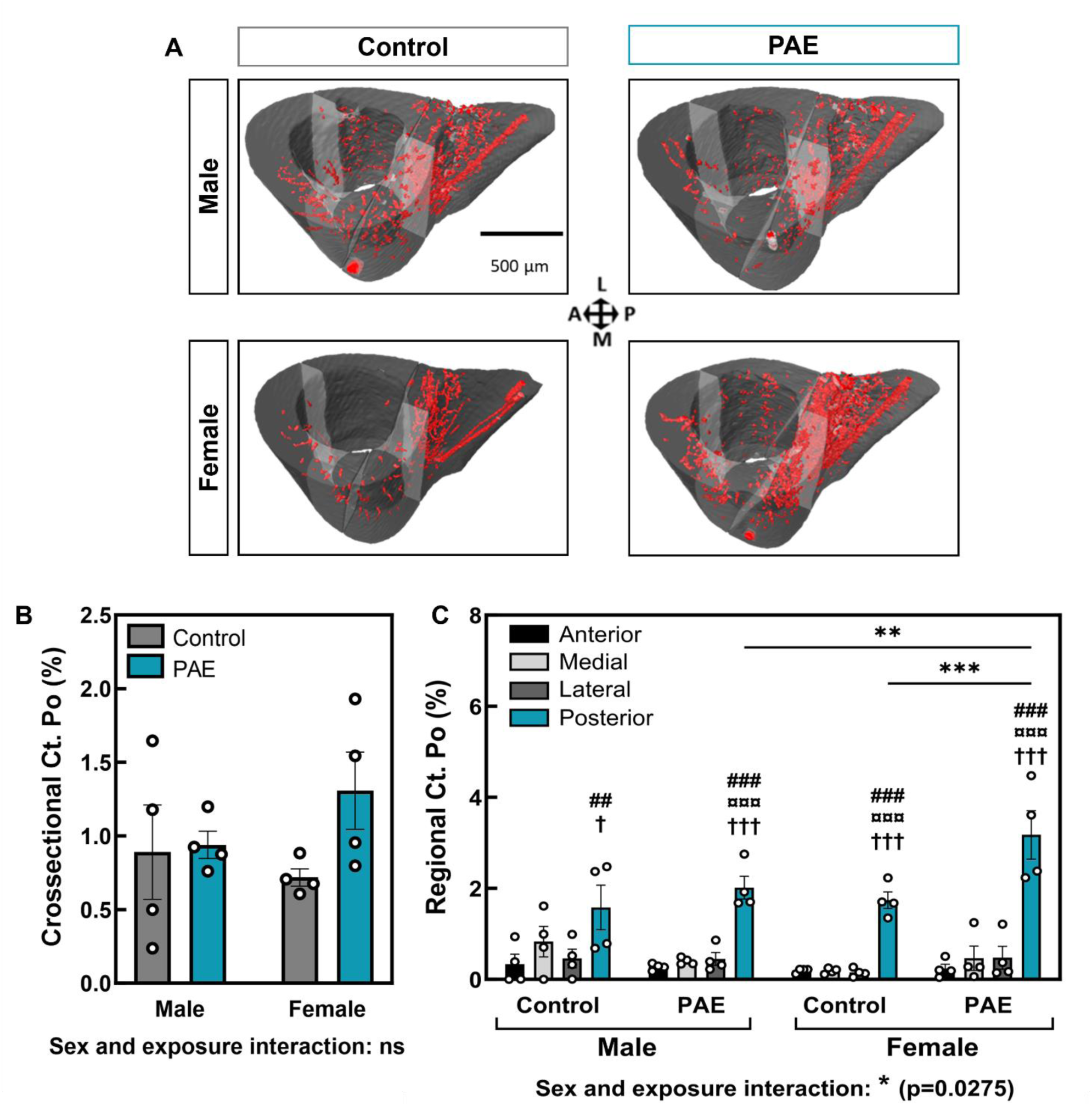
PAE females display reduced cortical porosity in the posterior region of the tibiofibular junction. (**A**) 3D renderings of intracortical canals (red) from male and female control and PAE mice. (**B**) Percentage cross-sectional cortical porosity and (**C**) percentage regional cortical porosity in anterior, medial, lateral and posterior regions. Data are presented as mean ± SEM with points showing individual animals. # denotes significant difference from anterior region within experimental group; ¤ denotes significant difference from medial region within experimental group; † denotes significant difference from lateral region within experimental group; * denotes significant difference between specific regions in different experimental groups. One symbol = p<0.05, two symbols = p<0.001, three symbols = p<0.001.

Three-point bending was performed to investigate whether the observed changes in bone microarchitecture and geometry with PAE influence bone material properties. Sex differences were observed in the control and PAE groups with maximum load (*p*<0.05), whereas stiffness was only reduced in control females compared to males (*p*<0.01; Table 2). Whilst no sex and exposure interaction was evident, PAE male and female mice showed a significantly decreased maximum load, and yield compared to control animals (*p*<0.05). However, a sex and exposure interaction (*p*=0.0489) was observed for stiffness and PAE male mice had significantly reduced bone stiffness compared to male controls (*p*<0.001; Table 2).

**Table 2.**
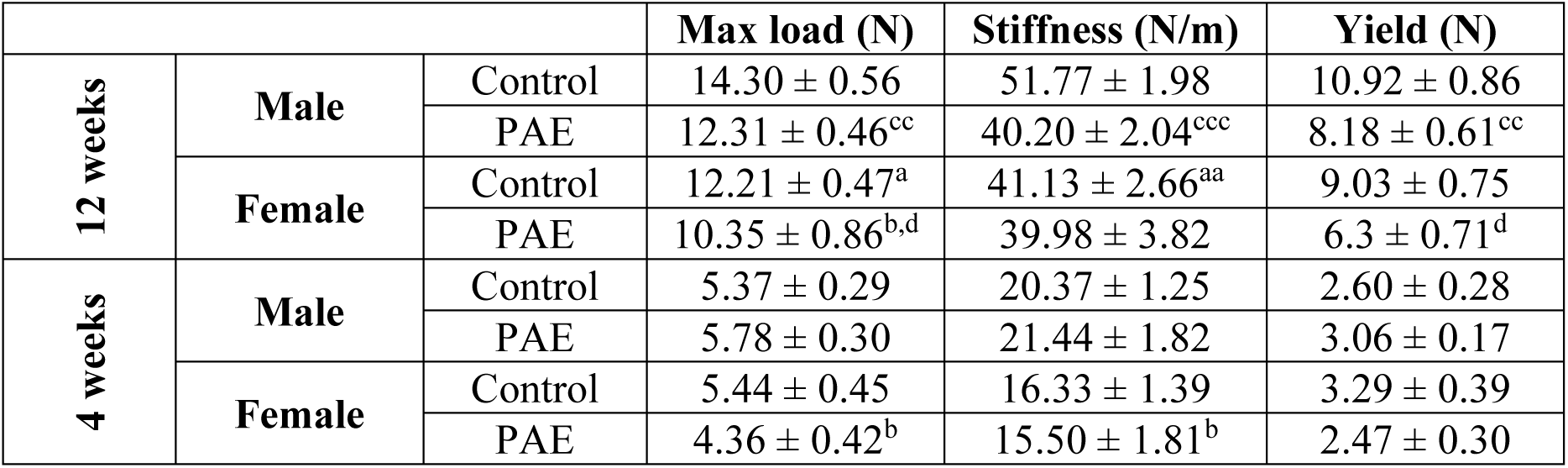
Three-point bending analysis of 12-week and 4-week-old control and PAE mice, in males and females. Data represented as mean ± SEM for n>5/experimental group. ^a^ = control male vs female, ^b^ = PAE male vs female, ^c^ = male control vs PAE, ^d^ = female control vs PAE. 1 symbol denotes p<0.05, 2 symbols denote p<0.01, 3 symbols denote p<0.001.

|  |  |  | Max load (N) | Stiffness (N/m) | Yield (N) |
| --- | --- | --- | --- | --- | --- |
| 12 weeks | Male | Control | 14.30 ± 0.56 | 51.77 ± 1.98 | 10.92 ± 0.86 |
| | | PAE | 12.31 ± 0.46<sup>cc</sup> | 40.20 ± 2.04<sup>ccc</sup> | 8.18 ± 0.61<sup>cc</sup> |
| | Female | Control | 12.21 ± 0.47<sup>a</sup> | 41.13 ± 2.66<sup>aa</sup> | 9.03 ± 0.75 |
| | | PAE | 10.35 ± 0.86<sup>b,d</sup> | 39.98 ± 3.82 | 6.3 ± 0.71<sup>d</sup> |
| 4 weeks | Male | Control | 5.37 ± 0.29 | 20.37 ± 1.25 | 2.60 ± 0.28 |
| | | PAE | 5.78 ± 0.30 | 21.44 ± 1.82 | 3.06 ± 0.17 |
| | Female | Control | 5.44 ± 0.45 | 16.33 ± 1.39 | 3.29 ± 0.39 |
| | | PAE | 4.36 ± 0.42<sup>b</sup> | 15.50 ± 1.81<sup>b</sup> | 2.47 ± 0.30 |

### PAE decreases growth plate width in males, despite no effects on bone length or growth plate bridging in either sex

At 12 weeks of age, females weighed less and had shorter tibias than males in both control and PAE groups (*p*<0.05; Table 3).

**Table 3.**
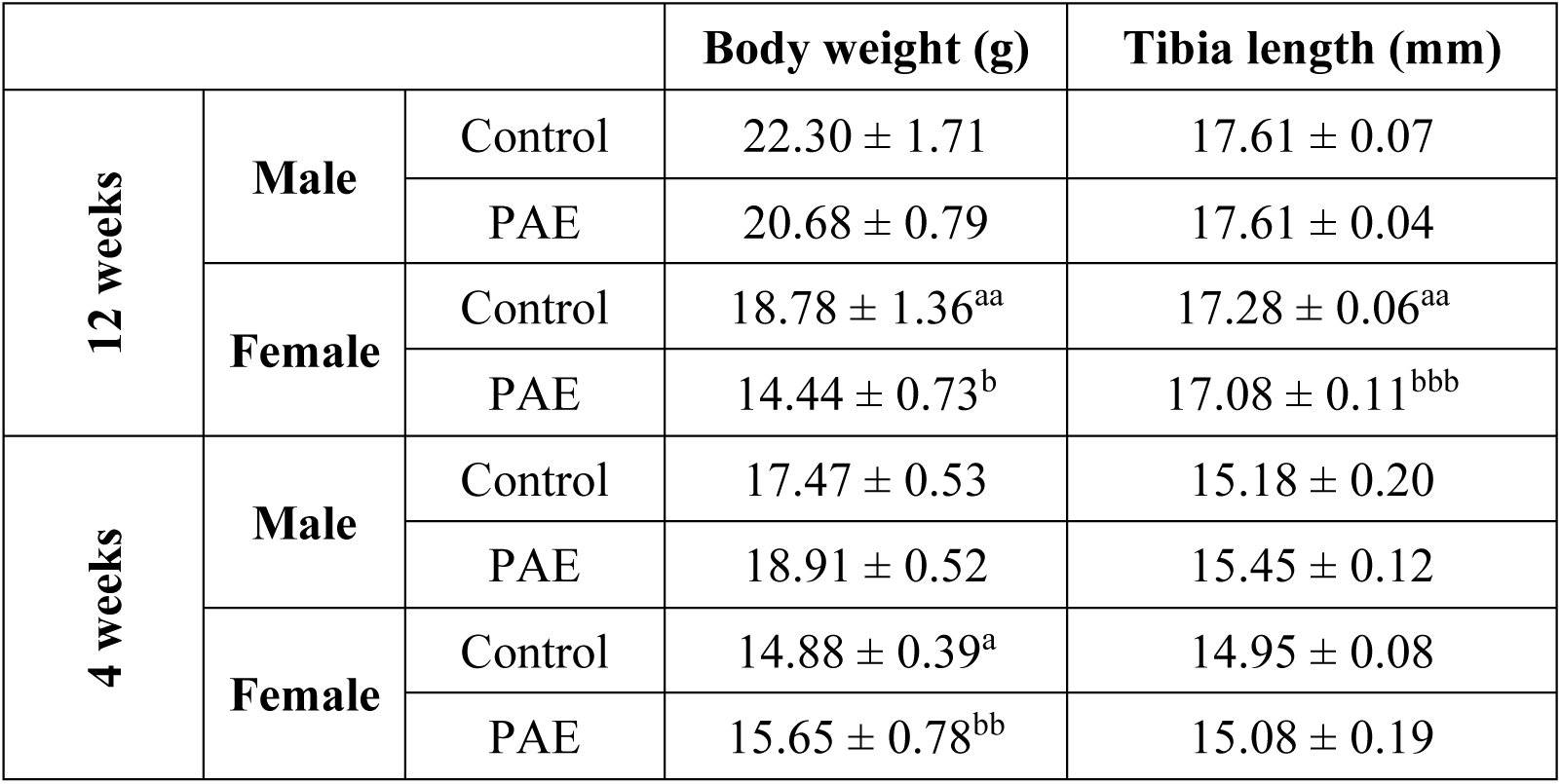
Weight and bone lengths of 12-week and 4-week control and PAE mice, in males and females. Tibial bone lengths were measured using µCT. Data represented as mean ± SEM for n>5/experimental group. ^a^ = control male vs female, ^b^ = PAE male vs female. 1 symbol denotes p<0.05, 2 symbols denote p<0.01, 3 symbols denote p<0.001.

|  |  |  | Body weight (g) | Tibia length (mm) |
| --- | --- | --- | --- | --- |
| 12 weeks | Male | Control | 22.30 ± 1.71 | 17.61 ± 0.07 |
| | | PAE | 20.68 ± 0.79 | 17.61 ± 0.04 |
| | Female | Control | 18.78 ± 1.36<sup>aa</sup> | 17.28 ± 0.06<sup>aa</sup> |
| | | PAE | 14.44 ± 0.73<sup>b</sup> | 17.08 ± 0.11<sup>bbb</sup> |
| 4 weeks | Male | Control | 17.47 ± 0.53 | 15.18 ± 0.20 |
| | | PAE | 18.91 ± 0.52 | 15.45 ± 0.12 |
| | Female | Control | 14.88 ± 0.39<sup>a</sup> | 14.95 ± 0.08 |
| | | PAE | 15.65 ± 0.78<sup>bb</sup> | 15.08 ± 0.19 |

Despite this, we observed no changes in total growth plate width with sex in either control or PAE groups (Fig. 6A & B). In both sexes, PAE had no effect on total body weight, or on tibial bone length, and no sex and exposure interaction was observed (Table 3). However, total growth plate width in male PAE mice compared to control was reduced (*p*<0.05), whilst female PAE mice compared to control were unaffected (Fig. 6A & B). Sex differences in the proliferative and hypertrophic zones of the growth plate were not observed in the control group, whereas an increase in the hypertrophic zone was seen in females compared to males in the PAE group only (*p*<0.05; Fig 6C & D). This was coupled with trends towards a reduction in the width of the proliferative zone, observed in both males and females with PAE compared to control (Fig. 6C), and in the hypertrophic zone (Fig. 6D). No apparent differences in TRAP labelling for osteoclasts were observed between males and females of either group or between control and PAE mice of either sex (Fig. 6A). Growth plate bridging assessment in these animals revealed a reduction in the number of bridges in female PAE mice compared to male PAE (*p*<0.001), whilst sex differences in areal density were observed in both control and PAE groups (*p*<0.01; Fig 6E-G). No significant differences were observed between control and PAE mice of either sex, but a significant sex and exposure interaction (*p*=0.0121) suggests there may be an effect on growth plate fusion (Fig. 6E-G).

**Figure 6.**
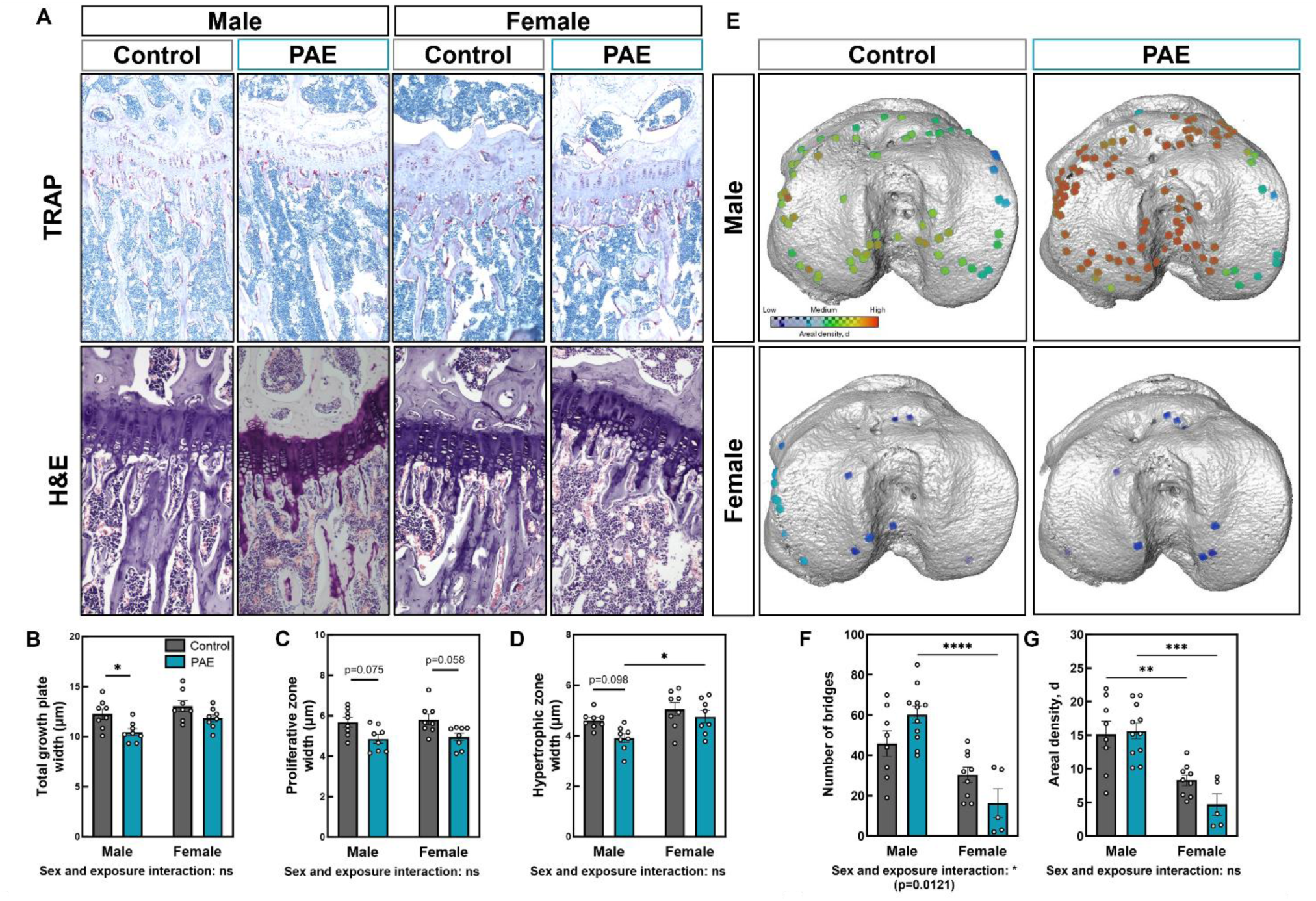
PAE decreases growth plate width in male mice. (**A**) TRAP and haemotoxylin and eosin-stained sections of male and female PAE and control 12-week-old mice. (**B**) Total growth plate zone width, (**C**) Proliferative zone width, and (**D**) Hypertrophic zone width, of male and female control and PAE mice. Ten measurements per section were obtained along the length of the tibial growth plate in the middle region of the knee joint. (**E**) Location and areal densities of bridges across the growth plate projected on the tibial joint surface in male and female control and PAE mice. **(F)** Number of bridges. **(G)** Areal density (d) of bridges defined as the number of bridges per 256 mm x 256 mm window. Data are presented as mean ± SEM with points showing individual animals. *= p<0.05, * p<0.01, ***= p<0.001.

### Juvenile mice do not display the altered skeletal phenotype with PAE that are observed in skeletally mature male and female mice

To assess whether the changes in the skeletons of PAE male and female mice are seen throughout development, we next investigated juvenile mice that are not yet skeletally mature. Whilst tibial lengths were unchanged between males and females, females weighed less than males in both the control and the PAE groups (*p*<0.05; Table 3). Body weight and bone length were unchanged in PAE mice compared to control in either sex (Table 3). Asides from tissue volume (Fig. 7A), similarly to 12-week animals, males and females differed across several trabecular parameters in both control and PAE groups (*p*<0.05; Fig. 7B-H). Interestingly, however, no sex and exposure interaction was evident and the detrimental effects of PAE to the trabecular bone microarchitecture observed in male 12-week-old animals were not observed at 4-weeks of age (Fig. 7 A-H, Suppl. Fig. 2). Similarly, no or limited effects on cortical bone geometry were observed in either sex with PAE compared to control, or in control males compared to females (Fig. 7I-N). However, sex differences in the PAE groups were observed, with PAE females displaying a reduction in cortical parameters along the tibial length compared to PAE males (*p*<0.05; Fig. 7I-N). This effect was also evident in bone strength measurements as no changes in biomechanical properties were observed in control mice compared to PAE in either sex, or in males compared to females in the control group (Table 2). Load and stiffness were reduced in females compared to males in the PAE group (*p*<0.05), whilst yield was unchanged. There was no sex and exposure interaction for any of these parameters.

**Figure 7.**
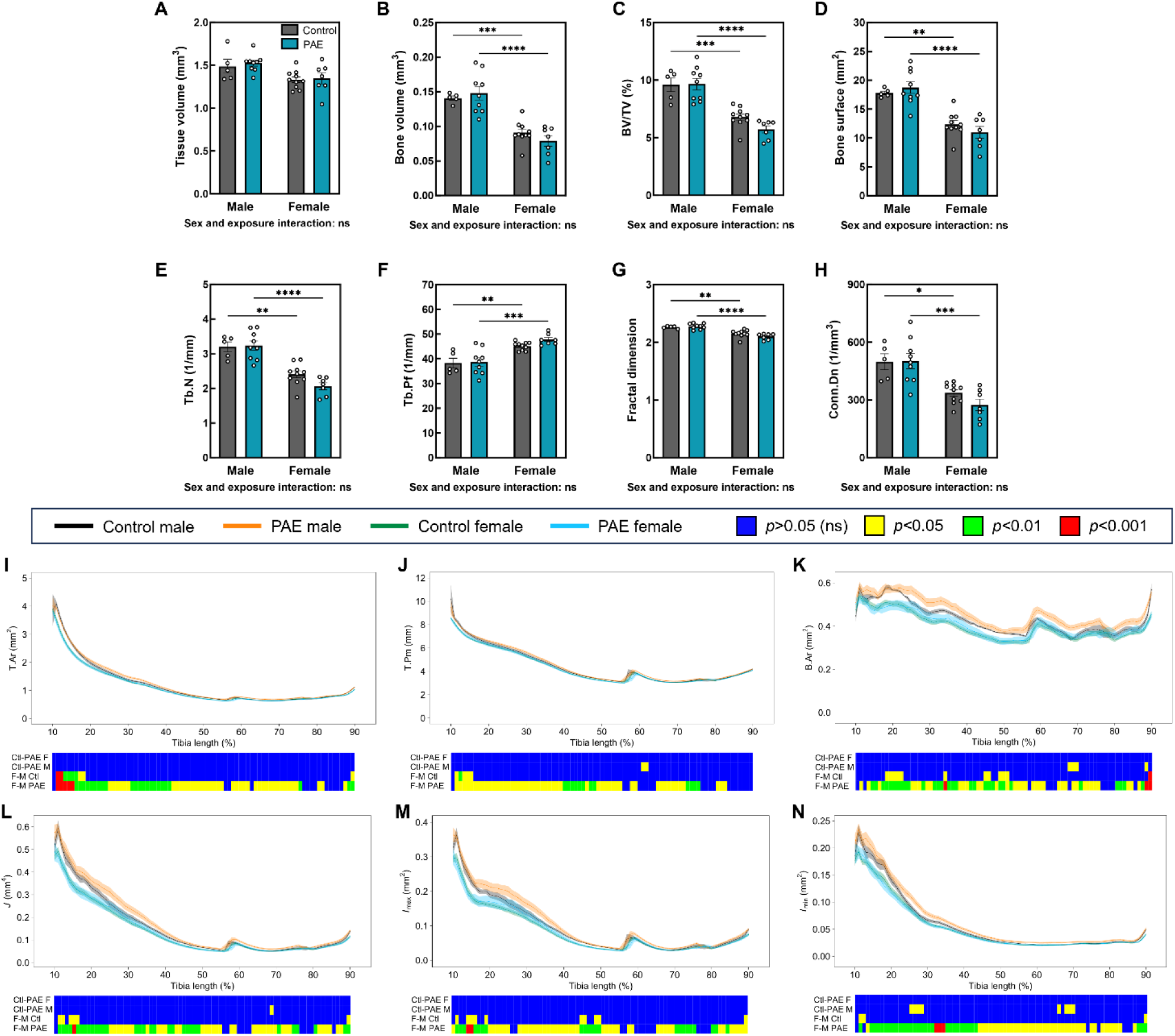
Juvenile mice do not display the altered skeletal phenotype that are observed in skeletally mature mice. µCT analysis of the trabecular compartment, including (**A**) tissue volume, (**B**) bone volume, (**C**) bone volume to tissue volume ratio (BV/TV), (**D**) bone surface, and (**E**) trabecular number (Tb.N), (**F**) trabecular pattern factor (Tb.Pf), (**G**) fractal dimension, and (**H**) connectivity density (Conn.Dn) in control and PAE males and females. Data are presented as mean ± SEM with points showing individual animals. *= p<0.05, ***= p<0.001, ****= p<0.0001. Measurement and statistical analysis heat map for (**I**) tissue area (T. Ar), (**J**) tissue perimeter (T.Pm), (**K**) Bone area (B.Ar), (**L**) resistance to torsion (J), (**M**) maximum second moments of inertia (I_max_) and (**N**) minimum second moments of inertia (I_min_) in male and female PAE and control mice. Line graphs represent mean ± SEM for male control (black), male PAE (orange), female control (green) and female PAE (blue) mice. Graphical heat map summarises statistical differences at specific matched locations along the tibial length (10% to 90%) for the PAE effect in females (Ctl-PAE F) and males (Ctl-PAE M), and the sex effect in control (F-M Ctl) and PAE (F-M PAE) groups. Red = p<0.001, green = p<0.01, yellow = p<0.05, blue = p>0.05 (not significant).

## Discussion

PAE has previously been shown to alter foetal skeletal development in animal models, however the long-term effects on the skeleton are unclear. Herein, we investigated the role of PAE on osteoblast function and gene expression in cells from male and female mice. Primary calvarial osteoblast function revealed reductions in mineralised nodule formation and sex-dependent changes on gene expression between control and PAE mice. To assess whether these alterations had any dimorphic effects later in life we conducted skeletal phenotyping of a murine model of PAE in both sexes, examining bones from skeletally mature and juvenile mice in the skeleton. In 12-week-old mice (skeletally mature), we observed clear differences between (i) sexes of each group and (ii) PAE and control animals within each sex in the growth plate, trabecular and cortical regions of the bone. However, how these differences presented were dependent on the sex. Whilst PAE male mice were affected along the entire length of the bone compared to control, PAE females only appeared to be affected in the cortical bone at the most distal end of the tibia and at the tibiofibular junction. Interestingly, 4-week-old juvenile mice did not possess these sex differences. This together suggests that the detrimental effects of PAE on the skeleton are sexually dimorphic (Fig. 8).

**Figure 8.**
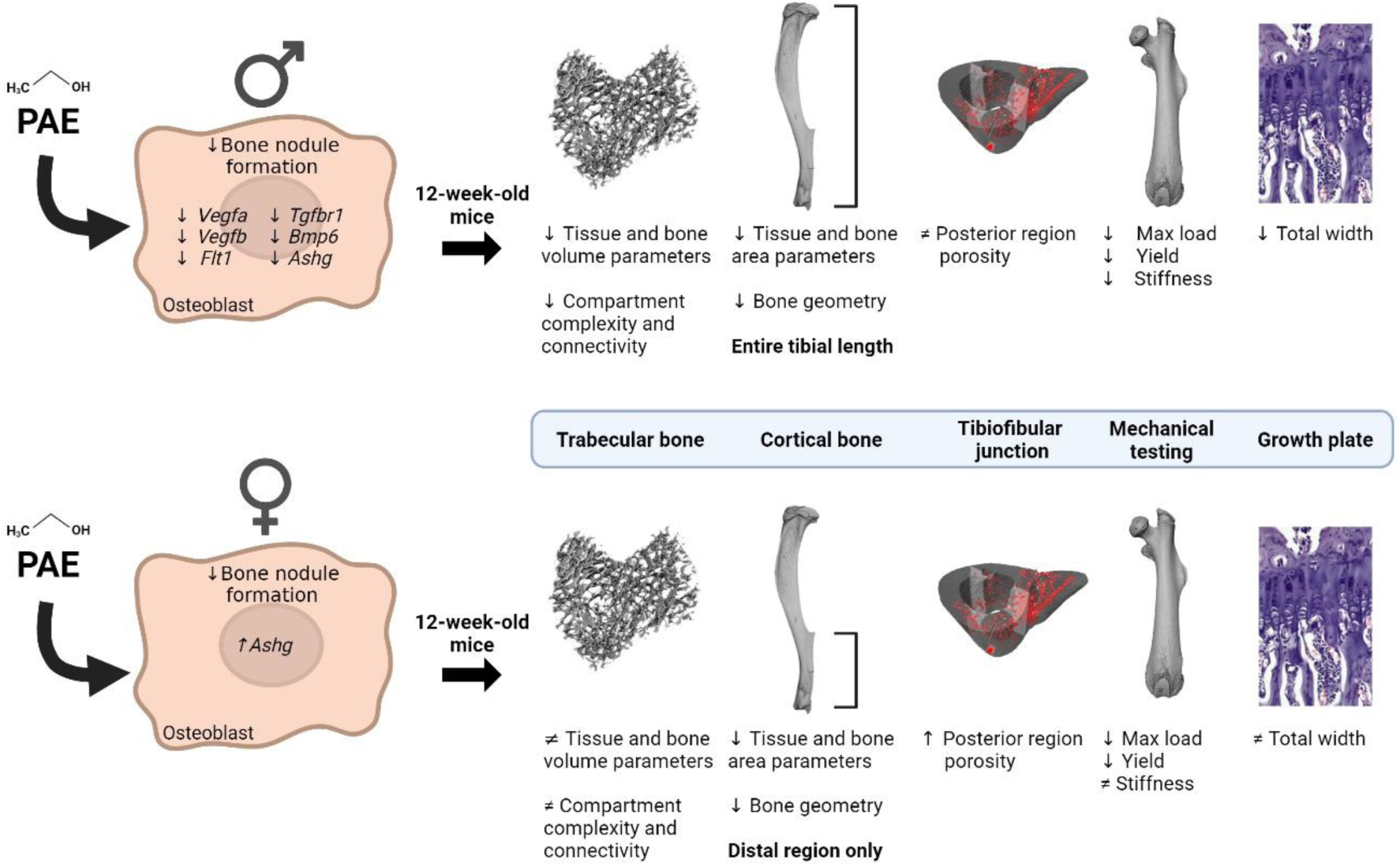
Sexually dimorphic effects of PAE on the skeleton. PAE osteoblasts from males and females produce less bone nodules than normal cells yet gene expression is differentially affected. This has lasting effects on the skeleton as PAE in 12-week-old male mice detrimentally alters the trabecular and cortical regions of the bone, bone mechanical properties and the growth plate. However, PAE female mice are protected in the growth plate and trabecular regions, but show detrimental effects only in the distal region of the tibia and at the tibiofibular junction.

The concept that the *in-utero* environment can alter the disease risk of a foetus later in life, termed as foetal programming, is widely accepted. The Barker hypothesis, in which foetuses that have adapted to limited nutrition during gestation are ‘programmed’ to be more at risk to cardiovascular diseases and diabetes in later life, gave an early indication that the prenatal environment can have lasting effects [49]. Disturbances to maternal diet and vitamin intake, smoking or substance abuse during pregnancy and the teratogenic effects of some chemicals and medications have all been shown to disturb foetal development [50–52]. Indeed, exposure to mycotoxins, environmental pollutants, medicinal products such as retinoic acid and methotrexate all result in skeletal abnormalities in rodent foetuses [53–55]. Further, limb defects have been noted in children prenatally exposed to cocaine and the deformities caused by thalidomide are well documented [56, 57]. Interestingly, long-term osteopenic effects of prenatal cocaine exposure have been observed in skeletally mature male rats at 32-weeks of age [58], whilst 5-month-old mice exposed *in utero* to the DNA methylation inhibitor, 5-aza-2-deoxycytidine, had negative alterations to their femoral microarchitecture e.g. bone volume fraction and trabecular number [59]. However, the long-term effects of compounds that do not have such severe or profound effects on the skeletal tissue has not been fully investigated.

Limiting alcohol consumption during pregnancy is now well accepted due to its negative impact on brain development, yet global prevalence of FASD remains high [11]. In human cohorts, the lasting effects of ethanol on the developing foetus, particularly in skeletal tissues, remain relatively unknown. This is in part due to the presence of confounding factors, including inaccurate self-reporting, polymorphisms in alcohol metabolism and coupling of alcohol with other insults e.g. smoking or illicit drug abuse [10, 60]. Animal models have the benefit of removing these confounding factors and controlling variables that human studies are limited by, alongside demonstrating dose-response relationships that can be employed to target different levels of PAE [60, 61]. Therefore, utilising preclinical animal models to investigate the sole effects of PAE (particularly low to moderate doses) on the skeleton is essential.

In this study, murine dams were given free access to drinking water containing 5% ethanol. No difference in water consumption was observed between control (water-only) and PAE-treated mothers, thus our model of PAE is more akin to long term, continual alcohol exposure that accumulates over time. A previous study demonstrated that the alcohol intake of breeding females in this murine model of 5% ethanol was ∼11g/kg/day, proportional to a human daily ethanol intake of approximately 100g, the equivalent of one bottle of wine [62]. Whilst consideration needs to be given to the sampling time as rodents primarily drink during the dark phase, leading to variability in blood alcohol concentrations, thus ultimately suggests that 11g/kg/day average-model used herein results in low-dose exposure [62]. This contrasts with other studies that look at the effects of binge drinking, single exposure or short-term exposure [26, 32]. For example, a single dose intraperitoneal injection of ethanol at gestational day 7 has been shown to increase prenatal mortality in mice and skeletal deformities, yet low-level exposure via ethanol inhalation over several days did not result in any abnormalities [26].

Interestingly, studies using animal models tend to show that male animals are more detrimentally affected by PAE than females [35, 62]. Indeed, individuals with FASD experience sex-related differences in clinical presentation, with males presenting significantly more neurodevelopmental impairment and females with more endocrine disorders [35]. Further, males with FASD exhibit larger volume reductions in brain structure than females [63]. However, human studies investigating FASD often do not report sex-specific differences either due to insufficient sample sizes or limited efforts to stratify for sex [64]. This is also true of studies specifically investigating skeletal effects of FASD. In their study on bone composition alterations in FASD children, Young and colleagues highlight the challenges surrounding combining male and female data and suggests that sex-specific effects may exist in this population [23]. This is to our knowledge, the first report of sex differences in the skeleton in pre-clinical models and therefore warrants further investigation of individuals with FASD to inform on stratified treatment.

Our initial analysis focused on calvarial osteoblasts isolated from neonates as this represents an effective model of *in vitro* bone formation [40]. We observed a clear reduction in bone nodule formation, regardless of sex with PAE, and our PCR array highlighted several genes that were differentially affected by PAE compared to controls in both males and females. Components of the TGFβ signalling pathway were downregulated in PAE osteoblasts from male mice, compared to control. This is consistent with previous studies which have shown that ethanol can affect TGFβ signalling in the brain [65–68] and the articular cartilage [69]. Defects in cleft palate formation are evident in FASD models and this process requires crosstalk between the TGFβ pathway and Sonic hedgehog (Shh) [70, 71]. Interestingly, the detrimental effects of PAE can be partially remedied by increased Shh [31], adding further evidence that these may be key pathways in the underlying pathology of FASD.

We also observed downregulation of genes associated with skeletal angiogenesis (*Vegfa, Vegfb and Flt1*). However, subsequent qPCR analysis confirmed this downregulation in PAE males during the differentiation stage of maturation (day 7), whereas the expression of *Vegfa* in female-derived osteoblasts was unchanged. Sexual dimorphisms in vascularisation and bone-derived VEGF signalling have previously been observed in a murine model [45]. Interestingly, bone-specific VEGF knockout altered parameters associated with cortical mass and geometry in females only, whereas despite elevations in cortical porosity in male mice, these measures were unaffected [45]. Given we did not observe alterations in VEGF expression in female-derived osteoblasts, this could explain the spatially limited effects of PAE on female bones at 12-weeks of age observed herein. However, the detrimental effects on the trabecular compartment in terms of bone mass and distribution of trabeculae observed in our skeletally mature male PAE mice contrasts with that seen in the bone-specific VEGF knockout. Furthermore, increased cortical porosity in the posterior region of the tibiofibular junction was observed in females, consistent with our previous findings [44], yet PAE females increased this further. In contrast, most interestingly, males were unaffected in this region. Evidence here suggests the complex nature of skeletal vascularisation, maintenance of bone and the sexually dimorphic actions of the genes involved, which could be affected by PAE.

Whilst we observed alterations in bone cell function and gene expression in cells derived from the calvaria, we did not examine the effects of PAE on the skull in our model. Other reports have noted craniofacial defects in murine foetuses at gestational day 17 [72, 73]. Irrespective of offspring sex, craniofacial malformations were observed equally in PAE, whereas body weight and length changes were only evident in PAE females [72]. Changes in skull size, shape variation and asymmetry have also been observed in pups at postnatal day 0, though this was not affected by sex [74]. Long-term assessment of craniofacial shape in rodent models, however, has not been performed and represents a possible avenue to further explore.

On examination of other areas of the skeleton, we observed alterations in growth plate width and number of bridges in male mice only, whilst females were unaffected. Despite this, neither group appeared to have altered growth, as demonstrated by no change in tibial length or body weight. This is in contrast to evidence that growth is impaired in PAE foetuses from rat and sheep models [29, 32, 34], but supports previous evidence indicating that in this murine model of PAE, weight changes in adult (3-6 months) were no different [62]. This may in part be explained as we did not perform longitudinal body length or weight assessment during skeletal development in these mice to assess whether growth deficiencies were initially present and subsequently compensated for or it could result from species differences. Given the trend towards increased growth plate bridging, suggestive of an accelerated fusion mechanism [48], this could explain the reasons for these results. Evidence of this from human populations is limited as FASD children aged < 9 years are similar in height to age-matched controls; by adolescence, individuals with FASD are shorter [23], yet longer term studies in skeletally mature FASD adults have not been performed. In addition, previous reports identified that adult female rats, and both juvenile and adult male rats exposed to alcohol *in utero* were more susceptible to developing an osteoarthritis-like phenotype and display cartilage degradation [69, 75].

Although we did not observe any immediately obvious damage to the cartilage or perform investigations into osteoarthritis in this model, given the effects of PAE on the underlying subchondral bone, further investigation is necessary to elucidate whether PAE has any effect on the progression of osteoarthritis, and whether this is the case in FASD adults.

Alterations in cortical composition and geometry were observed in PAE mice of both sexes, with male PAE mice detrimentally affected along the entire tibial length. Changes to cortical bone shape can be used to predict a reduction in bone strength, as a results of failing to resist excessive strains [43, 76]. Indeed, femurs from PAE males were weaker, less stiff and could sustain less load before undergoing plastic deformation than controls. In contrast, female PAE mice only displayed alterations in bone shape in the distal tibia and displayed reduced cortical porosity in the tibiofibular junction compared to controls. Whilst stiffness was unchanged, female PAE mice also showed reduced biomechanical properties (strength and yield). In a sheep model of PAE, similar reductions in tibial and femoral strength have also been observed, yet sex differences have not been reported [32, 33]. A study in FASD children has shown that they are more prone to fracture during early childhood, yet skeletal investigations in adults have not been performed [24]. Given these observations in preclinical models, further studies are warranted in human cohorts and to ensure that sex-stratification is included to delineate any potential sexual dimorphism.

Despite the profound effects of PAE at 12-weeks of age, we did not observe differences in PAE compared to control in either the males or females in our juvenile murine model (4-weeks of age), although some evidence of sex differences within the PAE group itself was evident. This therefore suggests that whilst there are inherent differences in the function and gene expression of primary osteoblasts derived from PAE neonates, the effect of PAE *in vivo* may require sex steroids and/or mechanical and environmental influences to exert its significant sexual dimorphic phenotype. Given that PAE leads to alterations in growth plate bridging, cortical geometry, and vascular changes at the tibiofibular junction, suggests that these changes may arise from altered responses to mechanical cues at these anatomical sites. Loading studies to assess the ability of these animals to respond to mechanical stimuli could be performed to further delineate this potential mechanism. Skeletal sex differences are observed post-puberty, demonstrating that sex steroids play a crucial role in maintaining life-long bone health. Our results suggest that PAE may affect men and women differently, independent of these sex hormones. Future studies aiming to understand how PAE affects the skeleton during sex hormone depletion could reveal novel mechanisms to aid sex-stratified treatments.

## Conclusions

In summary, evidence herein suggests that PAE has detrimental and sexually dimorphic effects on the skeletons of skeletally mature male and female mice. This effect is observed in spatially distinct regions of the bone and may be because of inherent differences in osteoblast function. These differences, however, were not evident in our juvenile model compared to controls and suggests a longer acting effect of PAE on the skeleton, coupled with distinct mechanoadaption between sexes when alcohol is given long term in low doses, which does not lead to immediately obvious clinical presentations. This study adds further evidence to indicate that the *in-utero* environment can have lasting impacts on skeletal health and that, for the first time, sex-specific changes in bone composition are evident with PAE, thus should be explored in human FASD populations to investigate this potential sexual dimorphism.

## Declarations

### Ethics approval and consent to participate

All experimental procedures were performed in accordance with the UK Animals (Scientific Procedures) Act of 1986 and regulations set by the UK Home Office and local institutional guidelines.

### Consent for publication

Not applicable

### Availability of data and materials

The datasets used and/or analysed during the current study are available from the corresponding author on reasonable request.

### Authors’ contributions

Study design: LEB, FMG, PG, NB, KAS; Analysis and interpretation of the data; all authors; Drafting the manuscript: LEB, KAS; Editing and approving final version: all authors.

## Acknowledgements

The authors would like to thank the Bioresources Unit at the University of Brighton for the assistance with the care of animal models in this study. We are also thankful to Dr Ruby Chang (Royal Veterinary College, UK) for her discussions regarding the statistical analysis. For the purpose of open access, the author has applied a CC-BY public copyright licence to any Author Accepted Manuscript version arising from this submission.

## Supplementary Information

**Suppl. Figure 1.**
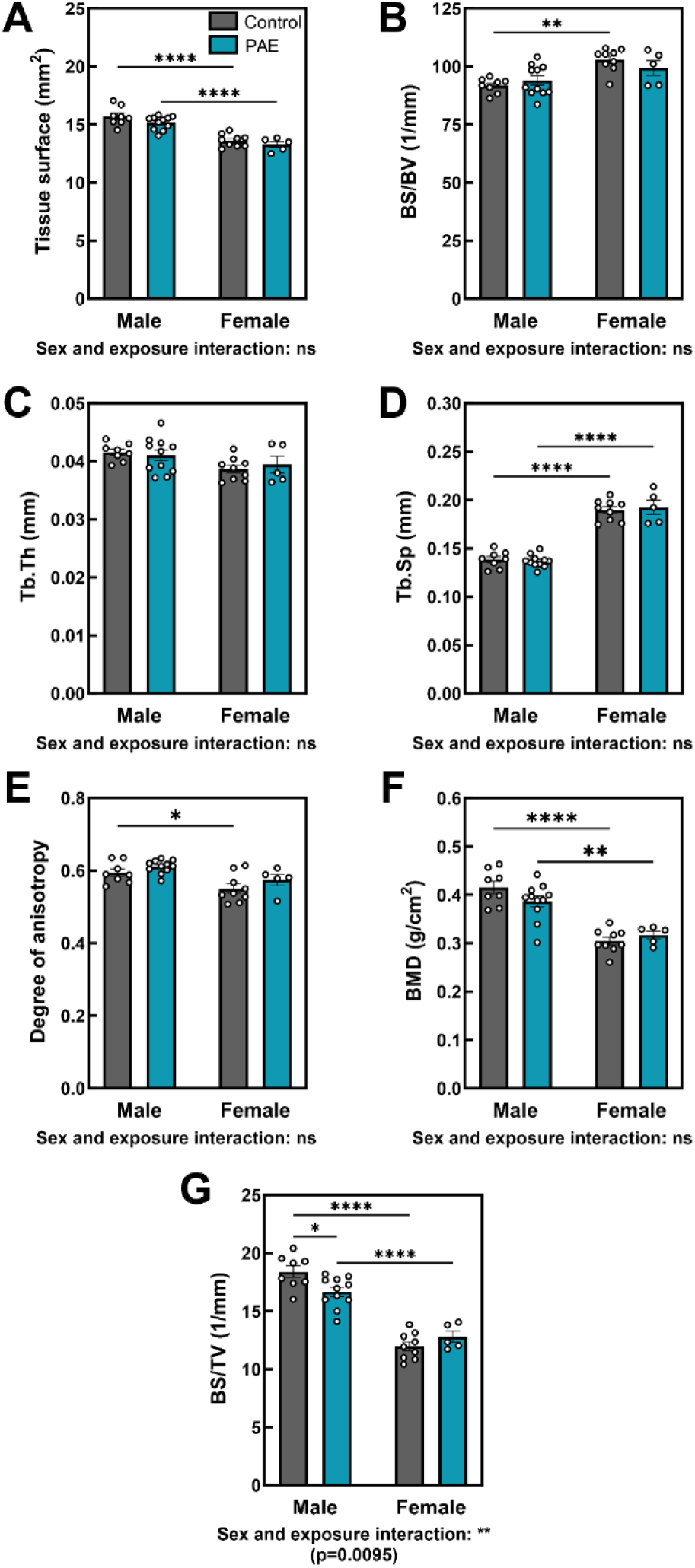
Additional trabecular parameters in 12-week-old mice. µCT analysis of trabecular bone parameters, including (**A**) tissue surface, (**B**) bone surface to bone volume ratio, (**C**) trabecular thickness, (**D**) trabecular separation, (**E**) degree of anisotropy, (**F**) bone mineral density (BMD), and (**G**) bone surface to tissue volume in PAE and control males and females. Data are presented as mean ± SEM with points showing individual animals. *= *p*<0.05, **= *p*<0.01, ***= *p*<0.001, ****= *p*<0.0001.

**Suppl. Figure 2.**
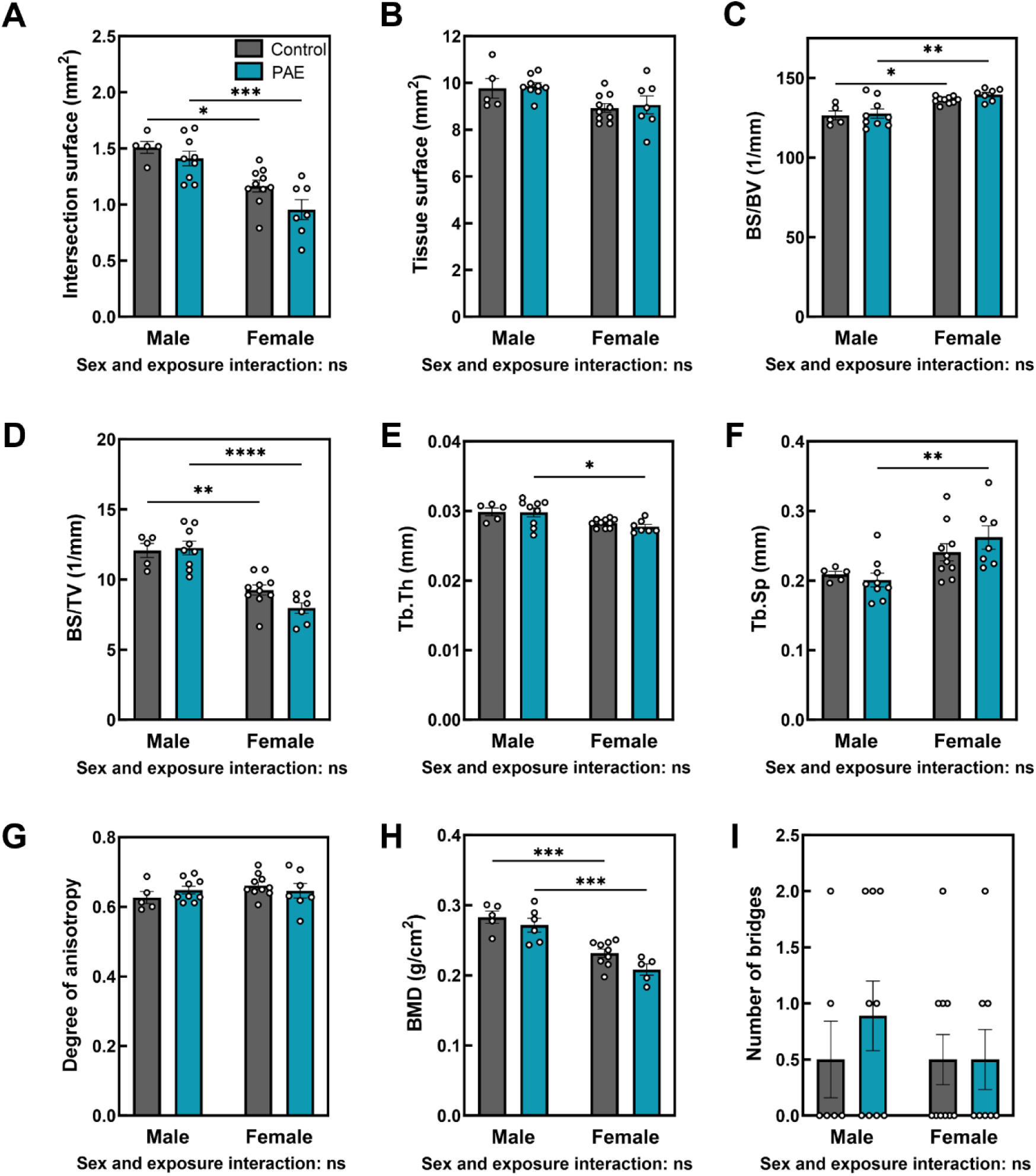
Additional trabecular bone parameters and growth plate bridges in 4-week-old mice. µCT analysis of trabecular bone parameters, including (**A**) intersection surface, (**B**) tissue surface, (**C**) bone surface to bone volume ratio, (**D**) bone surface to tissue volume ratio, (**E**) trabecular thickness, (**F**) trabecular separation, (**G**) degree of anisotropy, and (**H**) bone mineral density (BMD) in control and PAE males and females. (**I**) Number of growth plate bridges in control and PAE mice of both sexes. Data are presented as mean ± SEM with points showing individual animals. *= *p*<0.05, **= *p*<0.01, ***= *p*<0.001, ****= *p*<0.0001.

**Suppl. Table 1:**
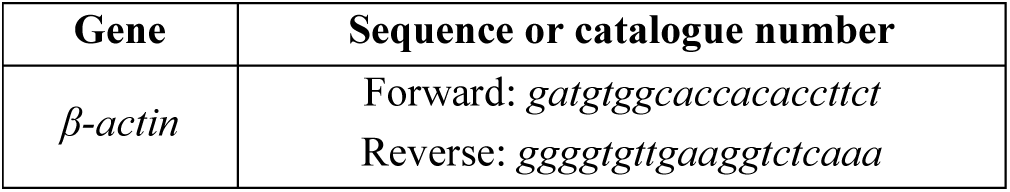
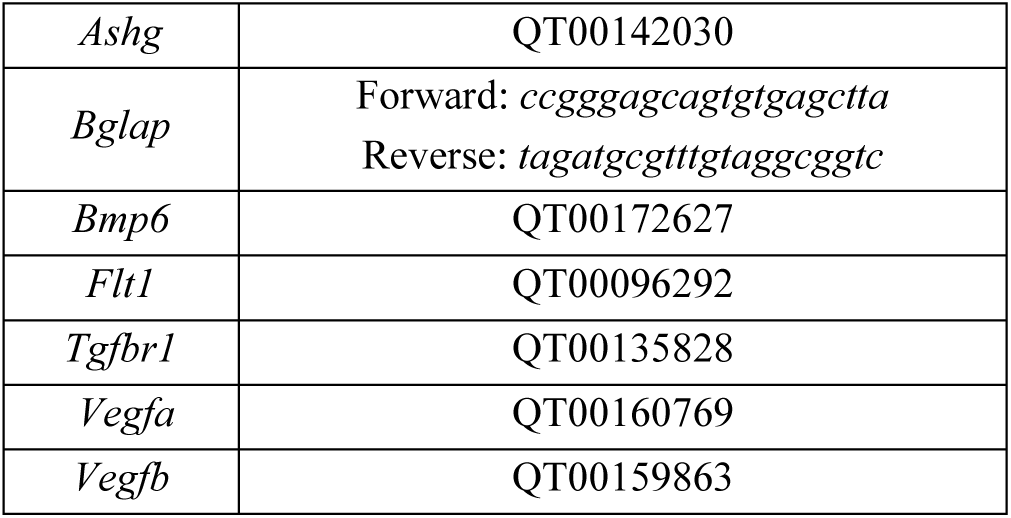
Primer sequences or catalogue numbers (QIAGEN QuantiTect primer assay) used for RT-qPCR.

